# A multi-omics view of wax synthesis in the wild cochineal bug, *Dactylopius opuntiae*

**DOI:** 10.64898/2026.08.13.744733

**Authors:** Julianne Dessert, Edward M. Marcotte, Howard Ochman

## Abstract

This study establishes a comprehensive genomic, transcriptomic, and proteomic foundation for the wild cochineal bug, *Dactylopius opuntiae*, a scale insect and agricultural pest of prickly pear cacti notable for its dense white waxy covering. Although fatty acyl reductases (FARs) are central to insect epicuticular wax biosynthesis, the genes underlying wax production in cochineal insects are largely unexplored due to a lack of genomic resources. We report a 359-Mb *de novo* genome assembly in which we identify 26 *FAR* genes. Through phylogenetic reconstruction of Coccoidea FAR enzymes we reveal lineage-specific expansions of tandemly arranged *D. opuntiae FAR* genes, and discuss these enzymes’ roles in the development of this insect’s epicuticular waxy coating.

## INTRODUCTION

The wild cochineal bug, *Dactylopius opuntiae* (Hemiptera), is an economically relevant scale insect due to its role as an agricultural pest of *Opuntia* prickly pear cacti (*1–3*). Like other cochineal species, *D. opuntiae* produces carmine, a brilliant-red pigment, although modern commercial supplies of the dye are primarily harvested from its cultivated congener, *D. coccus* (*3–6*). Cochineal scales are largely sessile insects that live on *Opuntia* cladodes (Fig. 1) and secrete a conspicuous white wax-like substance termed coccerin that covers their globular body (*7–9*). The cottony waxy coating of *D. opuntiae* is thicker and more voluminous than the powdery coating of *D. coccus* (*6*) and is thought to provide extra protection from predators, desiccation, and environmental toxins, and to help waft immature cochineal bugs to new plants (*1*, *3*, *6*, *9–12*).

**Figure 1.**
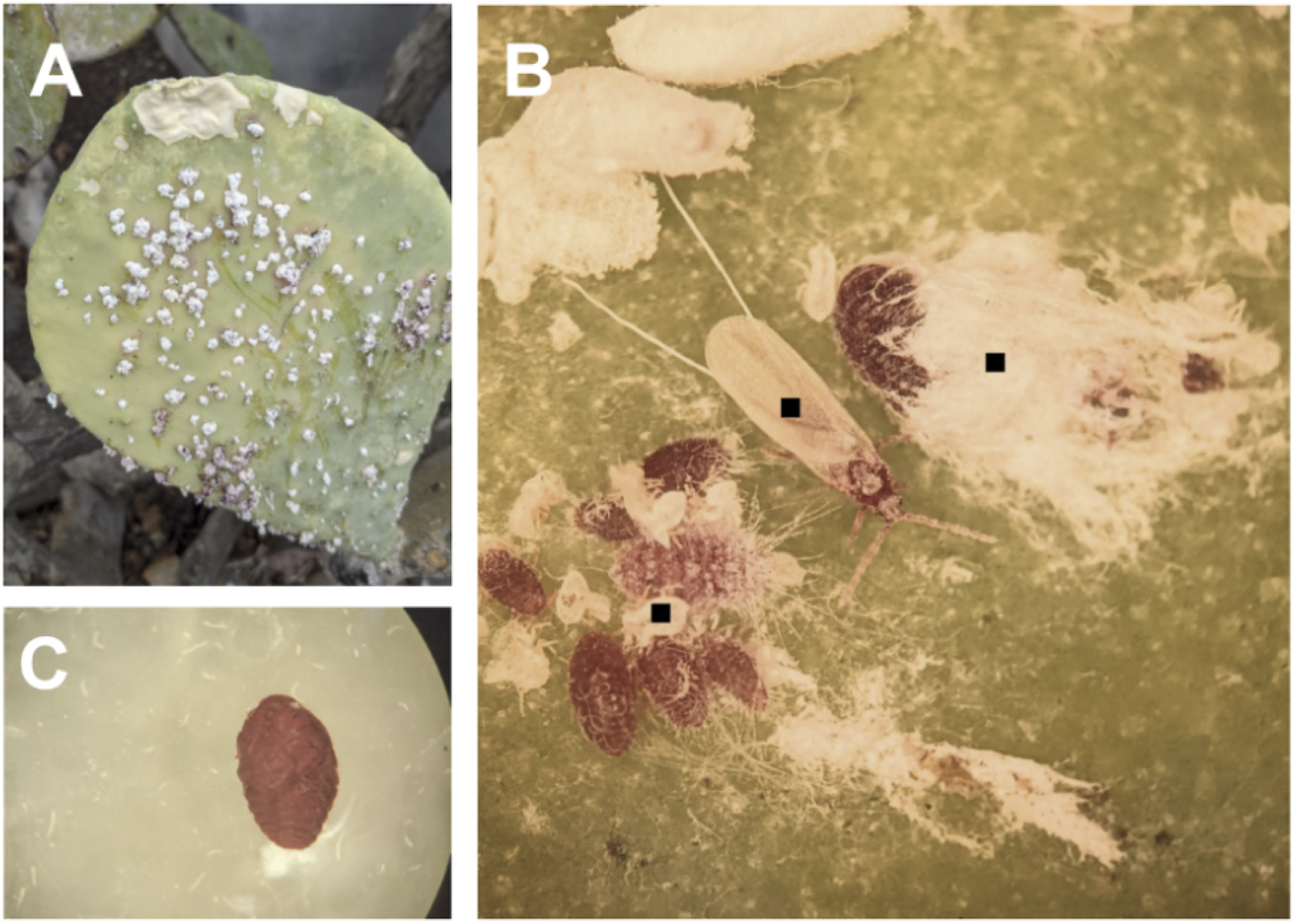
Habitat and life stages of the wild cochineal, *Dactylopius opuntiae*. (A) Cladode of *Opuntia* host cactus showing the density of white waxy coatings secreted by individual *D. opuntiae*. (B) Magnified view of *D. opuntiae* at different life stages. Black squares, from left to right, show a cluster of immatures excreting filamentous wax strands, an adult male with distinct slender and winged morphology, and an adult female insect almost entirely encapsulated by the waxy coating. (C) Isolated cochineal after treatment to remove the waxy layer revealing its carmine color and sessile morphological features.

Insect epicuticles comprise an assortment of lipid-based compounds—such as long-chain fatty acids, cuticular hydrocarbons (CHCs), wax esters, and fatty alcohols—with composition profiles varying across taxa (*11–14*). While there have been no analyses on *D. opuntiae’s* waxy coating, the primary components of *D. coccus’s* coccerin are CHCs and wax esters (*7*, *8*). CHCs are long-chain hydrocarbons and wax esters consist of a long-chain fatty acid (cocceric acid) and a fatty alcohol (cocceryl alcohol). The reduction of a fatty acid to its corresponding aldehyde and alcohol is catalyzed by a fatty acyl reductase (FAR) enzyme (*15*, *16*) and is a critical step for both CHC and wax ester production (Fig. 2). Following FAR activity, the metabolic pathway diverges—a fatty aldehyde can undergo oxidative decarbonylation to form a hydrocarbon (CHC) or a fatty alcohol and a fatty acid can be esterified to form a wax ester (*13*, *16–18*). The molecular biology underlying epicuticular phenotypes of scale insects is largely unexamined, and few insect FAR enzymes have been identified and experimentally characterized (*15*, *19–21*). Moreover, published cochineal genomes remain scarce and unannotated, precluding investigations into the synthesis of the insect’s signature epicuticular waxy coating.

**Figure 2.**
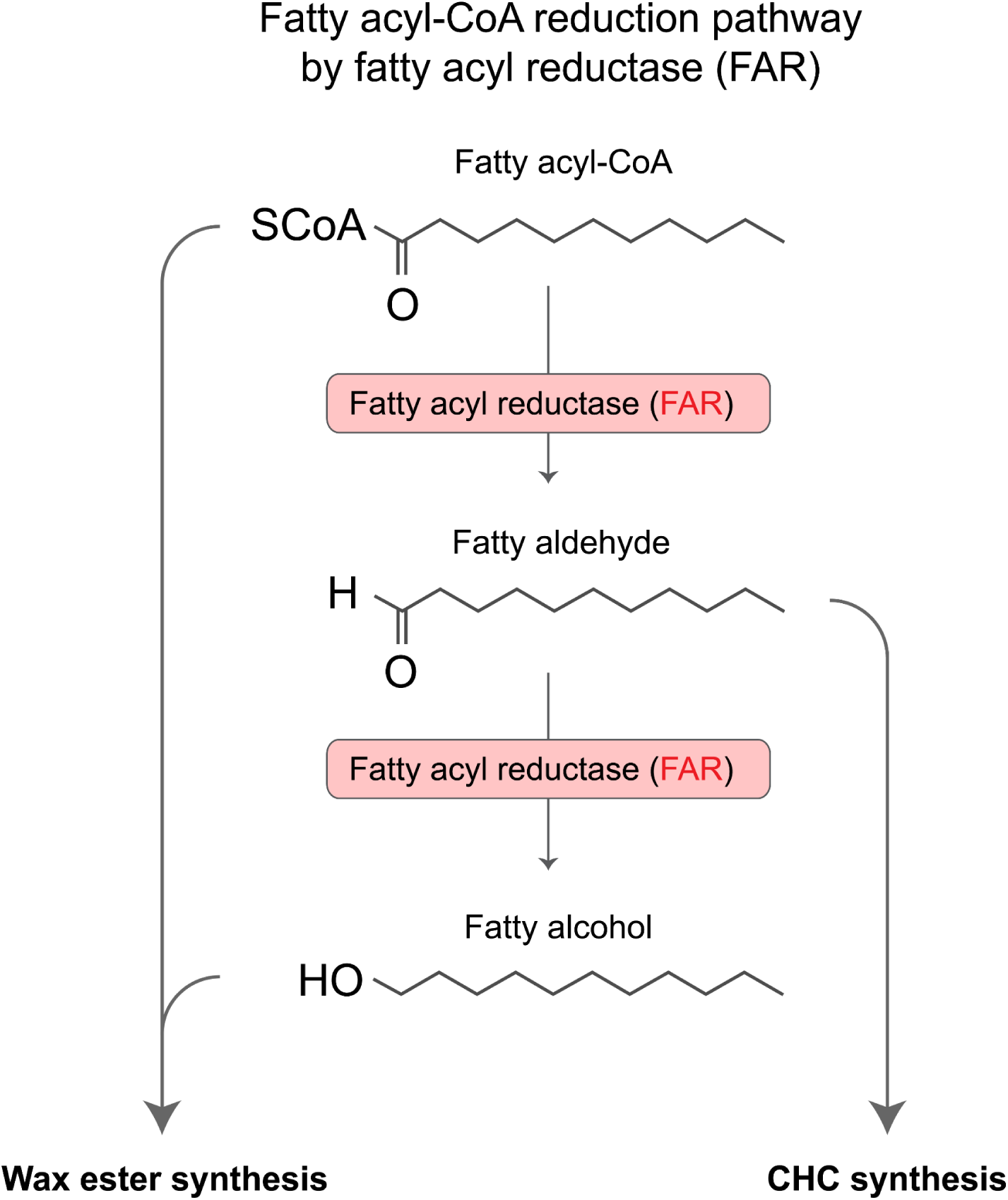
Pathway of fatty acyl-CoA reduction by fatty acyl reductase (FAR). FAR catalyzes two sequential reduction reactions required for epicuticular lipid biosynthesis. FAR first reduces fatty acyl-CoA to a fatty aldehyde, which is subsequently reduced by FAR to its corresponding fatty alcohol. Each of these three molecules serve as intermediates that are further processed by additional enzymes for either wax ester or cuticular hydrocarbon (CHC) synthesis. Fatty acyl-CoA molecules and fatty alcohols are esterified for wax ester synthesis, while fatty aldehydes are diverted for CHC synthesis.

*FAR* genes that are presumed to participate in epicuticle development have been reported in other Coccoidea insects, such as the Chinese wax scale and the cotton mealybug (*13*, *15*). Beyond their involvement in CHC and wax ester formation, FAR enzymes can also contribute precursors for pheromone production (*20*, *22*), regulate adult development and physiology (*23*), and determine female fertility (*23*). Given this diversity of functions, it is not surprising that insect *FAR*s constitute large multigene families arising from complex histories of duplications. For example, 17 *FAR* genes have been identified in *Drosophila melanogaster* (*24*), 26 in the Chinese wax scale, *Ericerus pela* (*25*), 36 in the pea aphid, *Acyrthosiphon pisum* (*24*), and 43 in the Asian citrus psyllid, *Diaphorina citri* (*24*). These expansions are striking in contrast to other organisms in that vertebrates, including humans, possess only two *FAR* genes, and some fungal species have none (*20*).

The underlying basis of the duplication and diversification of *FAR* genes in insects is beginning to emerge. A phylogenetic analysis of over 350 *FAR*s in 12 Hemipteran species revealed that while several duplication events occurred in the ancestors to the entire order, others are confined to certain lineages or species, and potentially defining specialized functions or epicuticular phenotypes (*24*). Here, we combine genomic, transcriptomic, and proteomic approaches to characterize the wild cochineal bug, investigate the patterns of *FAR* gene expansions in *D. opuntiae* and discuss how FAR enzymes may underlie development of its epicuticular waxy coating.

## RESULTS

### *D. opuntiae* genome assembly, gene prediction, and proteomic validation

We compiled a *de novo* genome assembly of the wild cochineal bug, *Dactylopius opuntiae,* using a combination of Nanopore long-read and Illumina short-read sequencing. Our genome assembly totals 359 Mb, comprises 1,474 scaffolds, and reports an N50 of 1.1 Mb and an average GC content of 31.2% (Table 1). The genome, transcriptome, and proteome completeness scores are all >91% based on the BUSCO Hemiptera odb10 gene-protein set. Gene prediction and subsequent longest isoform selection and removal of proteins shorter than 50 amino acids yielded 14,980 proteins. Of these, 9,766 (65.2%) proteins could be assigned to orthologous groups, with most annotations derived from Insecta (3,204 proteins). A total of 12,192 genes had a transcript abundance of >1 transcripts per million (TPM) (table S1). Additionally, peptides mapping to 2,138 of the predicted proteins were detected via mass spectrometry (MS), yielding a total of 17.5% of genes with confirmed transcript and protein expression (table S1).

**Table 1.** *D. opuntiae* genome assembly and annotation statistics.

| **Genome assembly and annotation** |  |
| --- | --- |
| **Assembly** |  |
| Total length (Mb) | 359.2 |
| Number of scaffolds | 1,474 |
| Scaffold N50 (Mb) | 1.1 |
| L50 | 104 |
| GC content (%) | 31.19% |
| **Masking** |  |
| Simple repeats (%) | 6.69% |
| **Gene Prediction** |  |
| Protein-coding genes | 14,980 |
| Total transcripts (isoforms) | 18,007 |
| Expressed genes (avg TPM > 5) | 9,015 (60.2%) |
| Proteins detected (Mass Spec) | 2,138 (14.3%) |
| **Annotation** |  |
| eggNOG ortholog hits | 9,766 (65.2%) |
| COG functional categories | 9,007 (60.1%) |
| GO term assignments | 6,335 (42.3%) |
| KEGG pathway hits | 5,759 (38.4%) |
| **Completeness** |  |
| Genome BUSCO | 91.40% |
| Transcriptome BUSCO | 91.90% |
| Proteome BUSCO | 92.40% |

Assembly continuity reported as N50 and L50 values. TPM (transcripts per million) is a measurement of transcript abundance. Completeness scores were assessed by BUSCO using the Hemiptera\_odb10 set.

### *FAR* genes in the cochineal genome

Reduction of fatty acids to their fatty alcohols by FAR enzymes is used in both wax ester and CHC synthesis (Fig. 2). Our cochineal genome contains 26 candidate *FAR* genes, which corresponded to 32 transcripts. FAR enzymes typically contain two conserved domains: an N-terminal Rossmann-fold NAD-binding domain and a C-terminal male-sterile domain (Fig. 3). Twenty-three of the 26 FAR proteins possessed both characteristic domains, with one additional FAR predicted to contain only the male-sterile domain. The remaining candidates lacking both domains probably represent truncated genes or gene fragments. MS experiments identified peptides mapping to five FAR proteins. Do-3 and Do-7 were the two most abundant, ranking in the 38^th^ and 49^th^ percentiles respectively, of the total observed proteins. Transcript abundance analysis ranked *Do-3* and *Do-7* in the top 47% and 92% of all transcribed genes (table S1).

**Figure 3.**
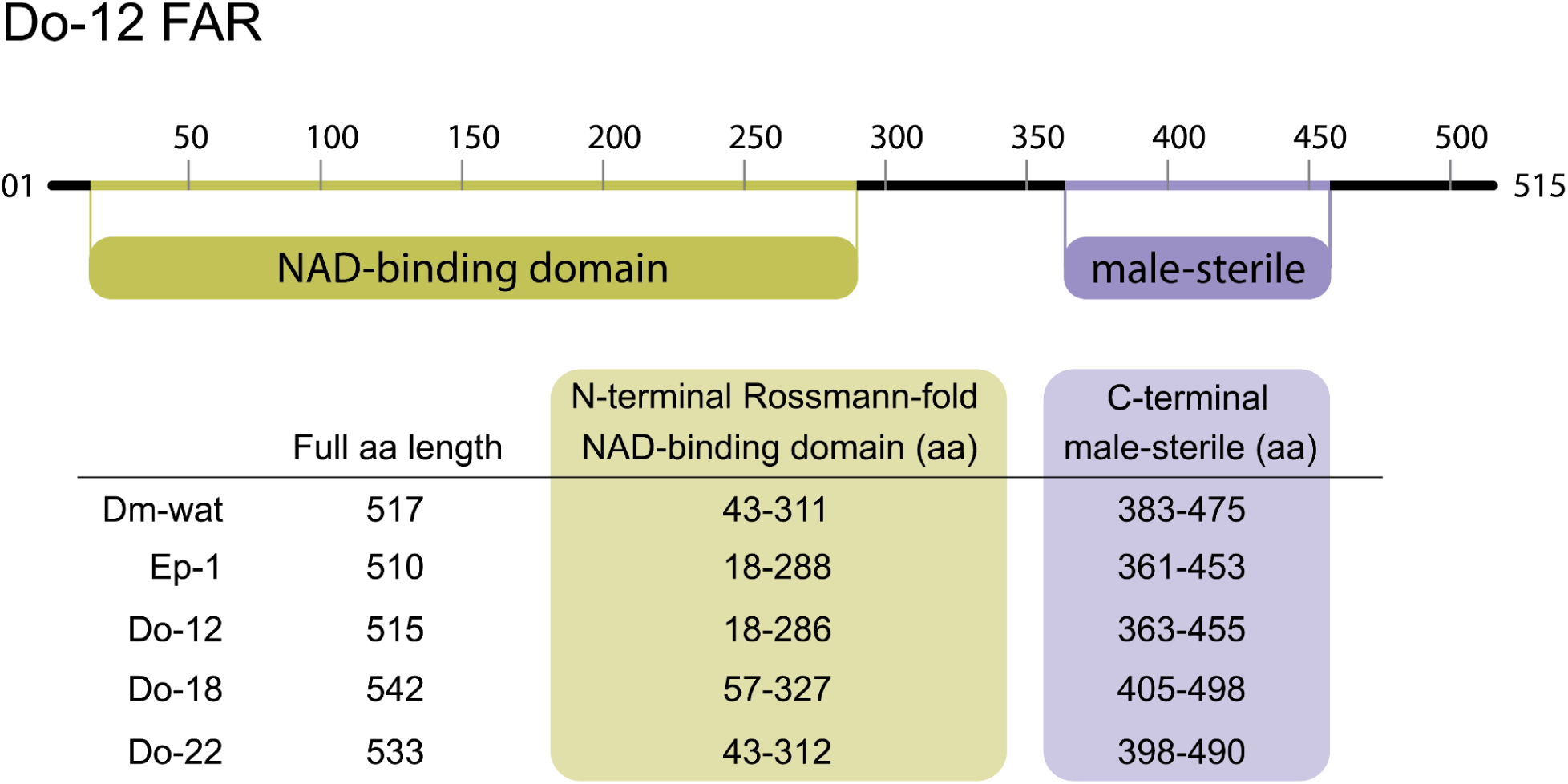
FAR proteins typically have two conserved domains. FARs typically possess both an N-terminal Rossmann-fold domain and a C-terminal male-sterile domain. Domains and locations of putative *D. opuntiae* FAR proteins were predicted using InterProScan (*59*). Dm-wat represents the characterized *waterproof FAR* gene product from *Drosophila melanogaster* (*19*) and Ep-1 refers to the FAR protein identified in the Chinese white wax scale insect, *Ericerus pela* (*16*). Do-12, Do-18, and Do-22 represent *D. opuntiae* FAR proteins identified in this study. Domain locations and linkage regions are consistent across all FAR proteins shown.

### Phylogenetic distribution and species-specific clustering of *D. opuntiae* FARs

Constructing a phylogenetic tree of the FAR proteins in *D. opuntiae,* other Hemiptera, and *Drosophila melanogaster,* places these *D. opuntiae* candidates within a narrow evolutionary context (Figs. 4 and 5). *D. opuntiae* FAR proteins are interspersed within the Hemipteran FAR clades, and all Hemipterans, including *D. opuntiae,* have FARs represented in multiple lineages. This distribution indicates that many expansions of the *FAR* gene family preceded speciation within the Coccoidea and prior to the divergence of the Dactylopiidae family that contains *D. opuntiae*.

**Figure 4.**
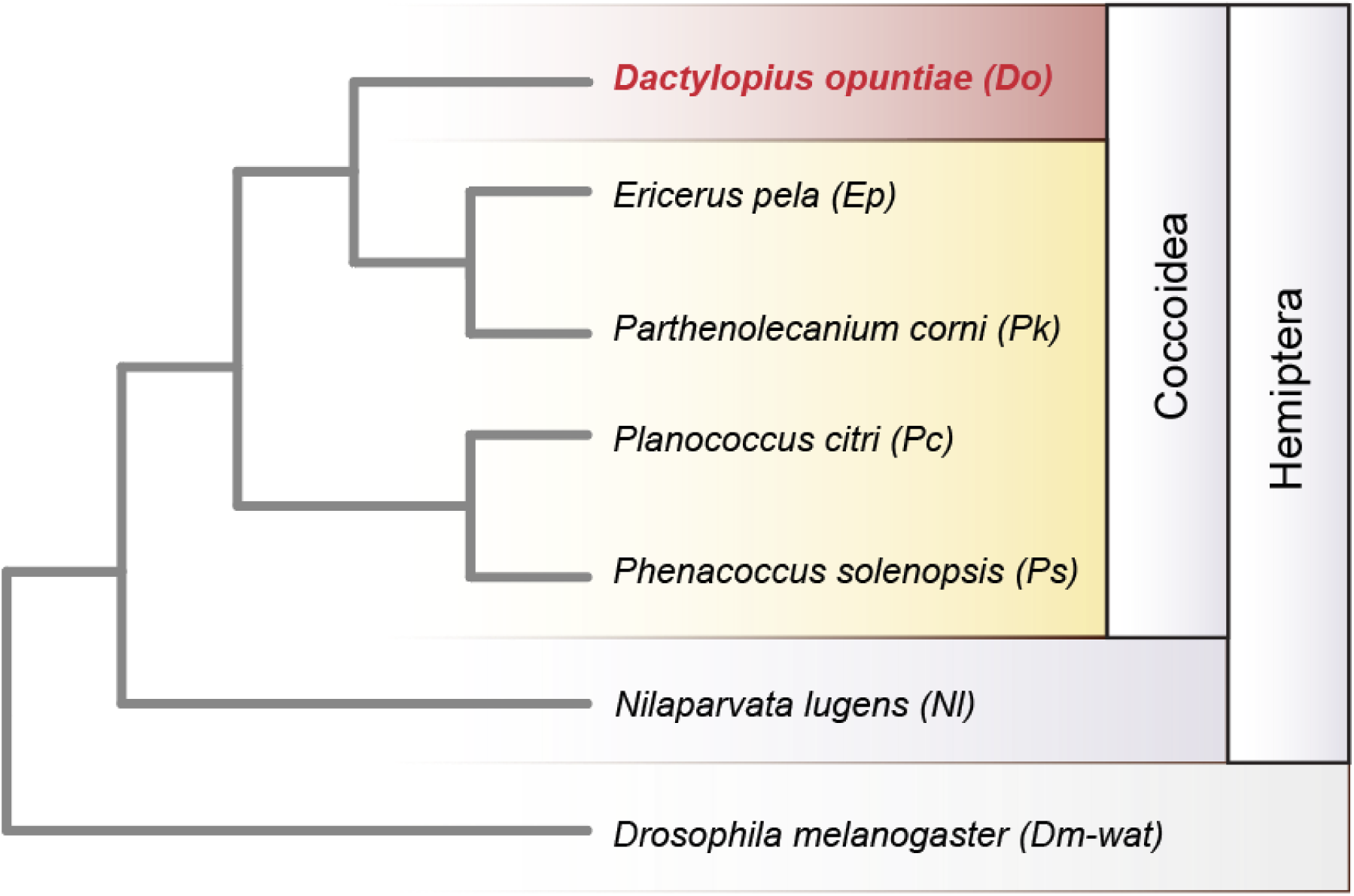
Species phylogeny of selected Coccoidea and outgroups. This species tree illustrates the phylogenetic relationships of scale insect superfamily (Coccoidea) members alongside outgroups *Nilaparvata lugens* (Hemiptera: Delphacidae) and *Drosophila melanogaster* (Diptera). *Dactylopius opuntiae* (*Do*) is the sole representative from the family Dactylopiidae. The two-letter species codes are used in the subsequent FAR protein phylogenetic analysis in Fig. 5.

**Figure 5.**
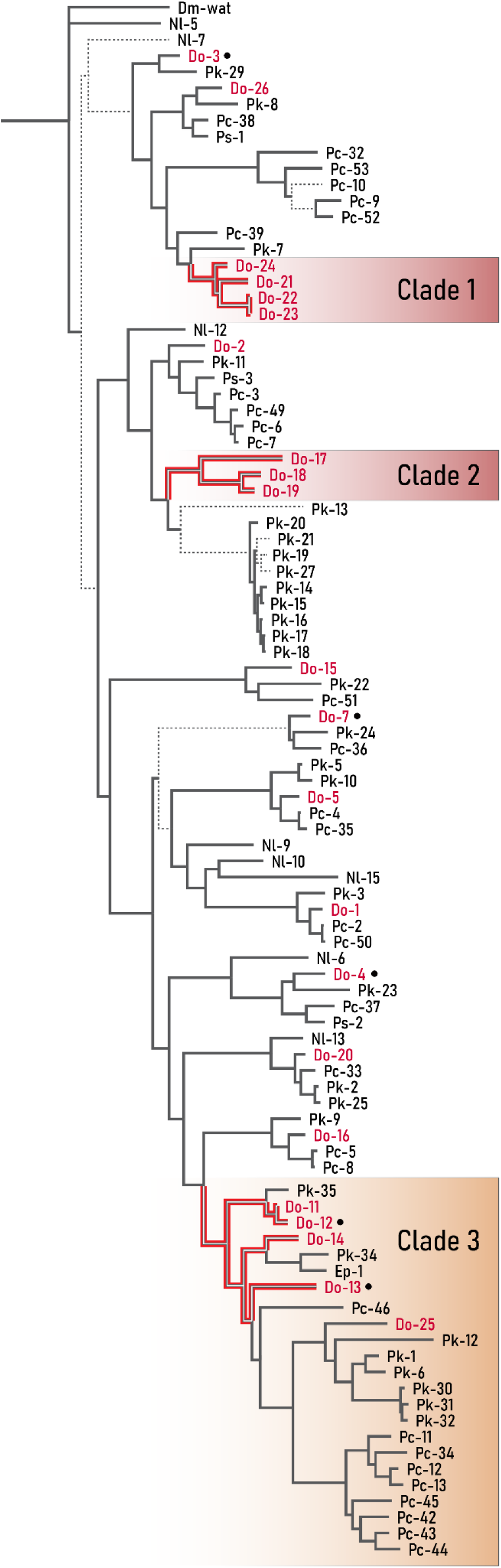
Diversification and lineage-specific expansions of FAR protein sequences in Coccoidea. **A** maximum likelihood tree was constructed from 96 FAR protein sequences pulled from seven insect species. Clades 1 and 2 highlight FAR clusters specific to *D. opuntiae*. Clade 3 indicates a paraphyletic group containing closely related *D. opuntiae* FARs as well as other Hemipteran FARs. Proteins labeled with black dots indicate those also identified by MS. Dotted lines denote branches with bootstrap support values below 75%.

In addition to these ancient events of gene duplication, there are two clusters of specific to *D. opuntiae*: Clade 1, comprising FAR enzymes Do-17, Do-18, Do-19, and Clade 2, comprising Do-21, Do-22, Do-23, Do-24, with their corresponding genes within each cluster arranged in tandem in genomic contigs 812 and 980, respectively. Proteins within each cluster span a wide range of protein sequence similarity: members of Clade 1 range from 48–80% pairwise sequence identity, and those within Clade 2 share 69–98% identity (Fig. 6). A third set of *D. opuntiae* tandem FARs sharing 38–91% amino acid identity sit in Clade 3 and exhibit a more complicated evolutionary history, with one pair (Do-11 and Do-12) arising from a recent duplication in *D. opuntiae*, and others (Do-13 and Do-14) that are more similar to FARs in other Hemipterans than to one another. MS data support the protein expression of two *D. opuntiae* FARs, Do-12 and Do-13, that reside in Clade 3.

**Figure 6.**
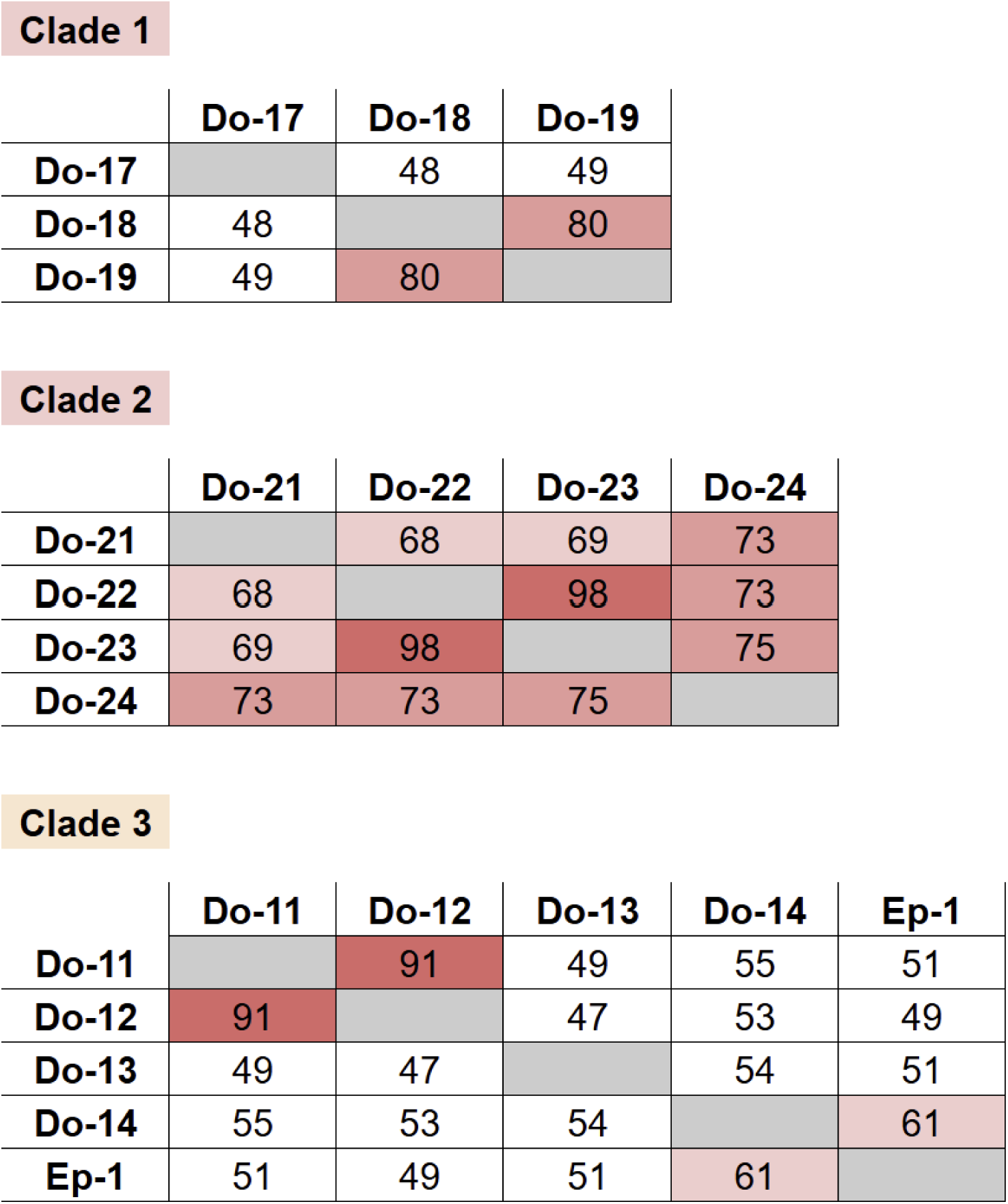
Protein sequence identities among *D. opuntiae* FAR clades. Percent amino acid identity between FAR orthologs within the three labeled clades in Fig. 5 are shown. Cells are shaded according to percent identity values with darker colors indicating greater similarity; higher identity values (e.g. >90% between Do-22 and Do-23 and between Do-11 and Do-12) indicate more recent gene duplication events.

### Functional inference of *D. opuntiae* FARs

Reference Hemipteran FARs, many of which have been experimentally characterized and are associated with wax synthesis, were included in our analyses to provide functional context. Some wax-related reference FARs cluster with *D. opuntiae* FARs, allowing functional inferences through phylogenetic proximity. First, Ep-1, resides in the paraphyletic Clade 3 and branches closest to Do-14 with 29% amino acid sequence identity. The *Ep-1* gene localizes to wax glands in male Chinese wax scales and is involved in synthesizing very-long-chain wax esters that create their white wax shield (*17*). Second, Ps-3, a P*. solenopsis* FAR branching off with Do-2 (63% sequence identity), generates cuticular hydrocarbons (CHCs) for the female’s cotton mealybug prolific waxy-filament phenotype (*13*). Third, another *P. solenopsis* FAR, Ps-1, clusters with Do-3 and Do-26 (47%, 64% sequence identity). Ps-1 is hypothesized to synthesize wax esters that form the base epicuticular layer across all life stages of the cotton mealybug (*26*). Alternatively, reference FARs that function in pathways not related to wax synthesis were included to identify FAR candidates involved in other functions. For example, Ps-2, implicated in pheromone synthesis in *P. solenopsis,* branches with Do-4 (60% sequence identity) (*26*). In summary, the topology demonstrates a widespread distribution of *D. opuntiae* FARs that span both ancient Hemipteran clades as well as more recent lineage-specific clusters.

## DISCUSSION

Our high-quality genome assembly of a Texas strain of *Dactylopius opuntiae* expands the very limited genomic resources currently available for cochineal scales (27). Our assembly integrates RNA-seq and MS data as evidence for gene prediction and functional annotation. Other cochineal assemblies include an unannotated chromosome-level assembly of *D. coccus* from Madeira, Portugal (GCA_977009245.1; ±360 Mb) (*28*) and an unannotated *D. opuntiae* assembly from a Chinese population (GWHFIWO00000000.1, ±326 Mb) (*27*, *29*). Notably, our ±359 Mb assembly is within 1 Mb of the *D. coccus* reference. While the assembly remains at a scaffold level rather than at a chromosomal scale, its high continuity and integration of transcriptomic and proteomic data are well-suited for subsequent gene identification.

Our identification of 26 *FAR* genes provides a foundational characterization of FAR enzymes in cochineal insects, offering key insights into the genes underlying cochineal wax synthesis. RNA-seq data confirming active expression of all but one of the predicted *FAR* genes provides evidence that these genes are neither assembly artifacts nor pseudogenes. Additionally, MS data confirms translation of five of these FAR enzymes, lending additional evidence that they are active components of the *D. opuntiae* proteome. The relatively high abundances of these FAR proteins underscore the metabolic investment *D. opuntiae* has made towards synthesizing both the FAR enzymes themselves and their downstream products. The *FAR* gene diversity in *D. opuntiae* is consistent with elevated numbers of *FAR* genes in other Hemipterans, supporting the hypothesis that the *FAR* gene family has expanded and undergone diversification to evolve multiple functions across the order.

A significant portion of these 26 *FAR* genes are likely involved in synthesizing the cochineal’s epicuticle, as suggested by several lines of evidence: First, as the FAR enzyme products are utilized in the production of a multitude of epicuticular-associated compounds (*14*), individual *FAR* genes could be tied to discrete metabolic pathways. Second, FAR enzymes can be highly substrate specific, restricted to catalyzing fatty acyl precursors of specific chain lengths, as shown in both *N. lugens* and *P. solenopsis* (*13, 23*). Third, epicuticle composition, along with *FAR* gene expression, is highly spatiotemporally regulated with transcript profiles varying across sexes, life-stages, and even tissues, as shown in *P. solenopsis* (*13*) and other studies (*17*, *23*, *26*). Although our transcriptomic and proteomic data are derived from whole-body rather than tissue-specific samples, precluding spatial localization of specific *FAR* genes to wax-secreting tissues, these patterns nevertheless suggest that an expanded *FAR* gene repertoire could help orchestrate the complex and dynamic epicuticle development throughout the insect’s life cycle. Beyond these epicuticular roles, we also expect that some of these *FAR* genes contribute to other biological processes, such as pheromone biosynthesis, as is documented in the silkworm (*20*).

To assess the timing of the *FAR* gene expansions, we analyzed the phylogenetic relationships of *D. opuntiae* FAR proteins to those in other Coccoidea species and included experimentally validated reference FARs to add functional context to our tree. This contrasts with previous Hemipteran FAR phylogenies which lacked strong representation from the Coccoidea superfamily (*24*, *26*). As demonstrated by the distribution of *D. opuntiae* FAR homologs in multiple lineages, epicuticular lipid synthesis does not appear to be governed by a single ancestral FAR, and instead involves multiple pathways. This distribution supports a Hemipteran *FAR* gene “birth-death” hypothesis (*24*), whereby the number of genes may increase via duplication and nonfunctional copies become inactivated or lost (*13*, *30*). For instance, the two *P. solenopsis* FAR paralogs (*Ps-1* and *Ps-3*), both involved in cuticular compound synthesis (*13*, *31*), occupy phylogenetically separate branches in the tree. Ep-1, which controls wax ester synthesis in the Chinese wax scale (*17*), branches off separately from other reference FARs that display similar functions. Recent phylogenetic analysis of Coccoidea genomes reports a radiation of scale insects alongside the diversification of angiosperms as the host plants (*27*). Based on the notion of multiple lineage-specific FAR diversifications, the absence of reference FARs in certain clades does not preclude involvement of the constituent *D. opuntiae* FARs in wax synthesis and rather presents them as candidates that may underlie unique cochineal biology.

Two lineage-specific *D. opuntiae* clusters represent recent tandem duplications and lack phylogenetic proximity to reference FARs. This genome architecture is notable, as tandem duplications have been implicated in the emergence of novel traits and adaptation of to environmental challenges (*32*, *33*). Accordingly, we hypothesize that because these *D. opuntiae FAR* genes are arranged in tandem, their enzymes may have evolved the capacity to synthesize diverse products (*32*, *33*). A third set of tandem *D. opuntiae* FARs in Clade 3 clusters with the reference Ep-1 from *E. pela,* suggesting these FARs function in synthesizing long-chain wax esters. The disparity between multiple *D. opuntiae* FARs with Ep-1 suggests the cochineal may have an expanded FAR toolkit for this function. Phylogenetic proximity to the Ep-1 reference FAR, coupled with our MS evidence, situates these Clade 3 *Do* FARs as strong candidates for the biosynthesis of wax esters comprising their waxy epicuticle. While these data suggest candidate FARs for wax synthesis, empirical validation—either from heterologous expression assays or *in-vivo* knockdown experiments—is ultimately required to confirm these candidates’ exact roles.

While recent progress has been made in sequencing and assembling scale insect genomes (*27*, *31*, *34*), there have been few attempts at functional annotation. Further expansion of genomic resources coupled with investigation into biosynthetic pathways can help resolve both taxonomic history and evolutionary timing of *FAR* gene expansions. Exemplified here in *D. opuntiae*, these data suggest that an expanded *FAR* gene repertoire shaped through iterative gene radiations has contributed to the diversity of unique epicuticular phenotypes observed across the superfamily.

## MATERIALS AND METHODS

### Insect collection and processing

Nymph and adult female wild cochineal bugs were harvested from cactus pads (*Opuntia sp.*) collected on the University of Texas at Austin campus and washed in 100% ethanol. To loosen and remove wax, insects were transferred to a 1:1 solution of xylene and 100% ethanol, shaken at 4°C for 12 min, and then washed in 100% ethanol at room temperature. Following treatment, insects were visually inspected, and any remaining wax or debris was removed with forceps under a dissecting scope. After removing wax, insects were washed with 70% ethanol, twice in DI water, and dried before flash-freezing in liquid nitrogen or placed at −80°C for later processing. Flash-frozen bugs were triturated by mortar and pestle, and stored at −80°C.

### DNA and RNA extraction

For Nanopore long-read sequencing, nucleic acids were extracted from 42 mg of powdered insects with the Quick-DNA Tissue/Insect Miniprep Kit (Zymo) except that after all reagents were added, samples were not subjected to bead beating in order to prevent DNA fragmentation. DNA purification was performed with the DNA Clean & Concentrator Kit (Zymo) and followed by RNase A (Thermo) treatment. DNA was concentrated using the DNA Clean & Concentrator Kit (Zymo), quantified, and stored at −80°C. For Illumina short-read sequencing, DNA was extracted from three frozen cochineal bugs with the QIAamp DNA Microkit (Qiagen), assessed for purity by Qubit and NanoDrop readings, and stored at −80°C.

Total RNA was isolated from 140 mg of flash-frozen powdered insects using the RNeasy Kit (Qiagen). Turbo DNase (Thermo) was added to disintegrate DNA, and RNA was purified using the RNA Clean & Concentrator Kit (Zymo). To selectively retain RNA fragments >200 nt in length, the binding step was adjusted to a 1:1:1 ratio (RNA:RNA binding buffer:ethanol). RNA integrity and quality were assessed by Nanodrop and on an Agilent Bioanalyzer.

### DNA and RNA Sequencing

DNA long-read libraries were prepared using a Nanopore Technologies SQK-LSK112 Ligation Sequencing Kit in conjunction with the NEB Compatibility Module (New England Biolabs), following the manufacturer’s instructions. Sequencing was performed on a Mk1B MinION with an R10.4.1 flow cell, producing a total of 4.81 million reads (±16 Gb), with 10.69 Gb passing QC (Phred score >8). For short-read sequencing, total DNA was pair-end sequenced on an Illumina Novaseq, generating a total of 353 million reads (±51 Gb), which were then quality-checked applying Q30-85%. Total RNA was subjected to polyA isolation, and the captured mRNAs were sequenced on an Illumina NovaseqXPlus, yielding a total of 328 million reads (±49 Gb).

### Genome assembly and annotation

DNA sequences generated from both Nanopore and Illumina platforms were trimmed and filtered using Trimmomatic (*35*), assembled with Flye (*36*), and used to polish with Pilon (*37*). The genome completeness was assessed with Quast (*38*) and BUSCO (v5.5.0; Hemiptera odb10) (*39*), repeats were masked using RepeatMasker within Funannotate (*40*, *41*), and the assembly was polished a second time with Pilon (*37*).

Raw RNA reads were trimmed and filtered with Trimmomatic (*35*) and mapped to the polished draft genome assembly using minimap2 (*42*). Gene prediction and annotation were performed with Braker3 (*43*, *44*) to generate a set of predicted protein-coding sequences. Functional annotations were assigned via ortholog assessment using eggNOG (*45*). Species identification to *D. opuntiae* was confirmed via BLAST analysis using mitochondrial cytochrome c oxidase subunit 1 (COI) barcodes retrieved from The Barcode of Life Data Systems (BOLD) (*46–48*).

### Protein preparation and mass spectrometry

Methods for preparation of samples for proteome analysis were adapted from Lin *et al.* (*49*) as follows: Powdered insects (7mg) were suspended in 50 μl cold PBS to which an equal volume of 4% SDS was added and incubated at 95°C for 10 min with shaking. Six volumes of acetone were added and incubation proceeded for 4 hr at 4°C, followed by centrifugation at 16,000 *g* for 15 min. Pellets were washed 2 times with acetone, air dried, resuspended in 100 μl solution of 1% sodium deoxycholate and 50 mM ammonium bicarbonate (HN_4_HCO_3_), and sonicated twice for 10 min. Samples were reduced in 5 mM TCEP at 56°C for 45 min and alkylated with 25 mM iodoacetamide in the dark at 22°C for 45 min. Solutions were quenched with 12 mM DTT and digested with trypsin overnight at 37°C. Reactions were terminated with 1% formic acid and centrifuged at 16,000 *g* to pellet precipitates. Supernatants were removed and filtered over Ultra 10 kd cutoff filter columns (Amicon), dried under vacuum and resuspended in 20 μl of Buffer A (0.1% Formic Acid, LC-MS water). Samples were desalted using 2 μg ZipTips (Millipore), and peptides were eluted in a solution of 0.1% TFA, 50% acetonitrile, 50% LC-MS water and dried under vacuum.

For MS/MS analysis, peptides were resuspended in 13 μl of Buffer C (0.1% Formic Acid, 5% acetonitrile, 95% LC-MS water) and analyzed on a Thermo Orbitrap Fusion Lumos Tribrid mass spectrometer. Peptides were separated by reverse-phase chromatography on a Dionex Ultimate 3000 RSLCnano UHPLC system (Thermo Scientific) with an Acclaim PepMap 100 C18 trap and EASY-Spray C18 reverse phase column (Dionex; Thermo Scientific) and eluted over a 3% to 40% gradient for 60 min. Data were collected using a data-dependent top-speed method with ions of charge R 2-6 selected for HCD and a stepped collision energy of 30%.

### Proteomic data analysis

Raw mass spectra were processed in MaxQuant (*50*) with the Andromeda search algorithm (*51*). Contaminating peptides were identified using a repository FASTA file compiled by Frankenfield *et al.* (*52*). Peptide spectral matches were analyzed against the cochineal proteome, and protein abundance values for peptides 6–30 amino acids in length were calculated using the iBAQ algorithm in MaxQuant (*50*). Results were filtered by applying a threshold of at least two unique peptides per predicted protein sequence with an FDR of 0.01. To quantify transcript and protein abundances, we reduced our gene and protein set by selecting the longest isoform per gene and excluding proteins shorter than 50 amino acids.

### Identifying candidate enzymes

A database of FAR proteins from various organisms was constructed to search for FAR homologs in the *D. opuntiae* proteome. FAR protein sequences were extracted from NCBI (*53*) and UniProt (*54*), and aligned in ClustalW (*55*). Top matches from three search methods—HMMER (*56*), MEME/MAST (*57*), and DIAMOND (*58*)—filtered by program default E-value cutoffs, were then used as BLASTP queries to further identify homology to other FARs. InterProScan (*59*) was used to identify Pfam protein domains in the FARs. FAR protein sequences used to build search algorithms are listed in table S2, and the resulting *D. opuntiae* FAR candidates are listed in table S3.

### Phylogenetic tree reconstruction

To place the *D. opuntiae* FARs in an evolutionary context, *FAR* genes from other insects, mostly Hemiptera (*Planococcus citri, Phenococcus solenopsis, Parthenolecanium corni, Nilaparvata lugens, Ericerus pela, Drosophila melanogaster*), were selected based on the following rationale: *FAR* genes from *N. lugens* and *E. pela* have been experimentally tested with documented diverse roles (*17*, *23*). *FAR*s from two mealybugs, *P. solenopsis* and *P. citri,* were included in order to supplement the limited genomic resources available for scale insects, and the *D. melanogaster waterproof* (*Dm-wat*) gene was included as the most well-studied insect *FAR* (*19*). *FAR* genes from *N. lugens* and *P. solenopsis* were obtained from published papers (*13*, *23*, *26*), while *FAR* gene sequences from *P. citri* (*60*) and *P. corni* (*61*) were extracted from published NCBI GenBank genomes by the procedure used for *D. opuntiae.* All *FAR* sequences were verified using BLAST against the NCBI non-redundant protein database before continuing. Genomes and FAR nucleotide and amino acid sequences utilized in this phylogenetic analysis are provided in table S4.

CDS sequences were conceptually translated using TranslatorX (*62*), and the amino acid sequences were aligned using MAFFT (*63*). Alignments were visually inspected and sequences shorter than 50% of consensus length were removed, leaving 22 of the 26 *D. opuntiae* FAR proteins. These remaining sequences were re-aligned and clipped using Clipkit (*64*) with the kpic-smart argument to preserve phylogenetically important regions. The final alignment of 96 FAR amino acid sequences was analyzed using iqtree2 (*65*) with ModelFinder (*66*). The maximum-likelihood tree was constructed using the best-fitting substitution model (LG+F+I+G4), and branch support was assessed with 1,000 ultrafast bootstrap replicates. The resulting tree was visualized with FigTree (*67*). Amino acid sequence identity matrices for each clade were calculated in Geneious (*68*).

## ACKNOWLEDGEMENTS

The authors gratefully acknowledge assistance from Dr. Nancy Moran, Dr. Zheng Li, Dr. Ophelia Papoulas, Tynan Gardner, and Emily Cook.

## AUTHOR CONTRIBUTIONS

Conceptualization: J.D., H.O., E.M.M.

Methodology: J.D.

Investigation: J.D.

Visualization: J.D.

Supervision: H.O., E.M.M.

Writing–original draft: J.D., H.O.

Writing–review & editing: J.D., E.M.M., H.O.

## DATA, CODE, AND MATERIALS AVAILABILITY

All data needed to evaluate the conclusions in the paper are present in the paper and/or the Supplementary Materials. Nucleotide data used in genome assembly, gene annotation, and protein prediction have been deposited in the NCBI database under the BioProject accession number PRJNA1490669. Proteomic data, including the raw mass spectrometry files, custom FASTA databases, and MaxQuant search parameter files have been deposited to the PRIDE/ProteomeXchange (*69*, *70*) database under project PXD080826. All data are accessible without restrictions.

